# *Myotis rufoniger* Genome Sequence and Analyses: *M. rufoniger*’s Genomic Feature and the Decreasing Effective Population Size of *Myotis* Bats

**DOI:** 10.1101/131904

**Authors:** Youngjune Bhak, Yeonsu Jeon, Sungwon Jeon, Oksung Chung, Sungwoong Jho, JeHoon Jun, Hak-Min Kim, Yongsoo Cho, Changhan Yoon, Seungwoo Lee, Jung-Hoon Kang, Jong-Deock Lim, Junghwa An, Yun Sung Cho, Doug-Young Ryu, Jong Bhak

**Affiliations:** The Genomics Institute, Ulsan National Institute of Science and Technology (UNIST), Ulsan 44919, Republic of Korea; Department of Biomedical Engineering, School of Life Sciences, Ulsan National Institute of Science and Technology (UNIST), Ulsan 44919, Republic of Korea; Personal Genomics Institute, Genome Research Foundation, Cheongju 28160, Republic of Korea; Geromics, Ulsan 44919, Republic of Korea; Department of Biomedical Science, School of Nano-Bioscience & chemical Engineering, Ulsan National Institute of Science and Technology (UNIST), Ulsan 44919, Republic of Korea; BK21 PLUS Program for Creative Veterinary Science Research, Research Institute for Veterinary Science, and College of Veterinary Medicine, Seoul National University, Seoul 08826, Republic of Korea; National Research Institute of Cultural Heritage, Cultural Heritage Administration, Daejeon 34122, Republic of Korea; Animal Resources Division, National Institute of Biological Resources, Incheon 22689, Republic of Korea

## Abstract

*Myotis rufoniger* is a vesper bat in the genus *Myotis*. Here we report the whole genome sequence and analyses of the *M. rufoniger*. We generated 124 Gb of short-read DNA sequences with an estimated genome size of 1.88 Gb at a sequencing depth of 66× fold. The sequences were aligned to *M. brandtii* bat reference genome at a mapping rate of 96.50% covering 95.71% coding sequence region at 10× coverage. The divergence time of *Myotis* bat family is estimated to be 11.5 million years, and the divergence time between *M. rufoniger* and its closest species *M. davidii* is estimated to be 10.4 million years. We found 1,239 function-altering *M. rufoniger* specific amino acid sequences from 929 genes compared to other *Myotis* bat and mammalian genomes. The functional enrichment test of the 929 genes detected amino acid changes in melanin associated *DCT*, *SLC45A2*, *TYRP1*, and *OCA2* genes possibly responsible for the *M. rufoniger*’s red fur color and a general coloration in *Myotis*. *N6AMT1* gene, associated with arsenic resistance, showed a high degree of function alteration in *M. rufoniger*. We further confirmed that *M. rufoniger* also has bat-specific sequences within *FSHB*, *GHR*, *IGF1R*, *TP53, MDM2*, *SLC45A2*, *RGS7BP*, *RHO*, *OPN1SW*, and *CNGB3* genes that have already been published to be related to bat’s reproduction, lifespan, flight, low vision, and echolocation. Additionally, our demographic history analysis found that the effective population size of *Myotis* clade has been consistently decreasing since ∼30k years ago. *M. rufoniger*’s effective population size was the lowest in *Myotis* bats, confirming its relatively low genetic diversity.

## Introduction

*M. rufoniger* is a species of vesper bat in the family Vespertilionidae [1]. It can be distinguished from other bats by its rusty orange fur (S1 Picture) and is therefore called ‘golden bat’ or ‘red bat’ in South Korea (Republic of Korea). Recently, its scientific name has been re-assigned to *M. rufoniger* from *Myotis formosus tsuensis* based on a molecular phylogeny and morphological study of bats [2]. Although its population is not assessed systematically, it is apparently a rare species that has only been collected in a handful of localities [2]. In South Korea, *M. rufoniger* is protected and designated as a natural monument. Being one of the most well-known and iconic protected wild animals in South Korea, even an *M. rufoniger* exhibition center is also in operation (golden bat exhibition center in Hampyeong County, South Korea).

In 2012, a fruit bat *Pteropus alecto* and an insectivorous *Myotis davidii* genomes were published, reporting bat-specific amino acid sequences on *TP53* (Tumor Protein P53) and *MDM2* (MDM2 Proto-Oncogene) genes that are associated with DNA damage checkpoint and DNA repair pathways. They provided insights into bats’ evolution of high metabolic rate and an increased amount of free radicals that are associated with flight [3]. In 2013, the Brandt’s bat (*Myotis brandtii*) genome analysis further identified unique amino acid sequence changes in *GHR* (Growth Hormone Receptor), *IGF1R* (Insulin Like Growth Factor 1 Receptor), *FSHB* (Follicle Stimulating Hormone Beta Subunit), *SLC45A2* (Solute carrier family 45, member 2), *RGS7BP* (Regulator of G-protein signaling 7 binding protein), *RHO* (Rhodopsin), *OPN1SW* (Opsin 1 [Cone Pigments], Short-Wave-Sensitive), and *CNGB3* (cyclic nucleotide gated channel beta 3) genes, providing new insights into bats’ delayed ovulation during hibernation, long lifespan, small body size, low vision, and echolocation [4].

As such, a set of close but distinct species genomes enable prediction of variants that may contain significant geno-phenotype association information [5-8]. This close species comparative genomics approach has been applied to *M. rufoniger* genome analyses to identify the species-specific variants that can confer functional and hence evolutionary adaptation of the *M. rufoniger*. To confirm and further identify such species-specific or bat-specific sequence changes, it is critical to evaluate such amino acid changes with as many genomes as possible at different levels of background comparison species. Thus, it is important to build biological resource with continuous sequencing of various species genomes.

A complete mitochondrial genome of *M. rufoniger* has been published already [9], while no whole-genome sequence has been reported yet, thus limiting the investigation of autosomal genetic signatures for environmental adaptations of *M. rufoniger*. Furthermore, the demographic history of a species can only be reconstructed accurately from deeply sequenced whole genomes [10,11]. Here, we provide a whole genome analysis of *M. rufoniger* by producing massively parallel short DNA sequences (approximately 124 Gb) with its genomic features and unique amino acid sequences, accompanied by its demographic history and genetic diversity.

## Results

### Whole genome sequences of *M. rufoniger*

The genomic DNA from the wild carcass of *M. rufoniger* found in Gosudonggul cave, Danyang, in South Korea, was sequenced using Illumina HiSeq2000 platform. A total of 124 Gb of paired-end short DNA sequences were produced with a read length of 100 bp, and target insert sizes of 566 bp and 574 bp from two genomic libraries. After reducing low sequencing quality reads and possible microbial contaminated reads, we acquired a total of 115 Gb of DNA sequences (Table 1; S1 Table). To confirm the species identification of the sample, a phylogenetic analysis was performed using a multiple sequence alignment of mitochondrial cytochrome b sequences, and our sample was verified to be the closest to *M. rufoniger* (S1 Figure). We performed a *K*-mer analysis (*K*□= □17) using the *M. rufoniger* whole genome sequences, and its genome size was predicted to be approximately 1.88 Gb (S2 Figure; S2 Table). This is similar to those of other bat species and smaller than those of other mammals [3,4,12]. Since there is no *de novo* assembled *M. rufoniger* genome, the DNA reads were aligned to all three available *Myotis* bat reference genomes (*M. brandtii*, *M. davidii*, and *M. lucifugus*; Table 1; S3 Table). The consensus DNA sequences of *M. rufoniger* were generated by substituting *M. rufoniger* single nucleotide variants (SNVs) detected from the whole genome sequencing data against the three reference genomes. The genome coverage (≥10×, 87.63 %) was the highest when the *M. lucifugus* assembly was the reference, while the coding sequence (CDS) coverage (≥10×, 96.60 %) was the highest when the *M. davidii* genome was the reference. The number of SNVs (58,660,193) was the lowest when the *M. brandtii* was the reference, while the number of small insertions and deletions was the lowest when the *M. daividii* was the reference. However, when the *M. brandtii* assembly was the reference, the mapping rate (96.50 %) and the number of consensus genes (17,247) was the highest. Therefore, we used *M. brandtii* reference-guided *M. rufoniger* consensus sequences for following analyses.

**Table 1.** Sequencing and mapping statistics of *M. rufoniger* genome.

|  | Myotis bat references |  |  |
| --- | --- | --- | --- |
|  | *M. brandtii* | *M. davidii* | *M. lucifugus* |
| Sequenced reads | 1,240,507,648 |  |  |
| Low quality filtered reads | 1,152,154,630 |  |  |
| Contamination filtered reads | 1,149,260,980 |  |  |
| Mapped reads | 1,109,045,915 | 1,101,733,443 | 1,095,435,354 |
| Mapping rate | 96.50% | 95.86% | 95.32% |
| Deduplicated reads | 1,023,540,144 | 1,017,982,320 | 1,010,388,456 |
| Depth coverage | 49.5 × | 51.5 × | 49.1 × |
| 1× genome coverage | 90.17% | 87.65% | 92.29% |
| 5× genome coverage | 87.83% | 85.68% | 89.88% |
| 8× genome coverage | 86.47% | 84.60% | 88.52% |
| 10× genome coverage | 85.58% | 83.88% | 87.63% |
| 1× CDS coverage | 98.86% | 99.10% | 98.13% |
| 5× CDS coverage | 97.68% | 98.19% | 96.29% |
| 8× CDS coverage | 96.60% | 97.33% | 94.88% |
| 10× CDS coverage | 95.71% | 96.60% | 93.81% |
| The number of homozygous SNVs | 50,615,560 | 56,635,858 | 50,441,069 |
| The number of heterozygous SNVs | 8,044,633 | 8,008,552 | 8,306,592 |
| The number of homozygous indels | 4,197,521 | 4,231,431 | 4,313,780 |
| The number of heterozygous indels | 517,253 | 457,868 | 512,018 |
| The number of consensus genes | 17,274 | 16,419 | 17,141 |
| The number of consensus genes without gap | 12,841 | 12,618 | 12,941 |

We examined the base composition of *M. rufoniger* genome (S4 Table) and found that the ratio of GC content was higher in the CDS region (52.15 ∼ 52.95 %) than that of the whole genome (41.49 ∼ 42.28 %), and this is concordant with the other mammalian species (the GC content ratios of the other species were 48.98 ∼ 54.35 % in the CDS region and 37.82 ∼ 42.93 % in the whole genome; S5 Table). The proportion of known transposon-derived repeats in the *M. rufoniger* genome was 19.05%, which is comparable with those of other bats (19.34 ∼ 33.32 %), but significantly lower than those of other mammalian genomes (25.27 ∼ 51.62 %; S6 Table).

The body of the *M. rufoniger* carcass used in this study was damaged and its morphological sex determination was not possible. We therefore compared its heterozygosity in X-chromosomal scaffolds with that of the other male *Myotis* bat (S7 Table). Since mammalian males have both X and Y sex chromosomes while females have two X chromosomes, a male individual has to show a lower heterozygosity in X chromosome than that of a female individual. Our sample showed an even lower heterozygosity in X-chromosomal scaffolds than that of the male *M. davidii* individual, indicating that it is clearly a male.

### Relationship of *M. rufoniger* to other mammalian species

We identified orthologous gene clusters from 13 mammalian genomes (seven bat genomes: *Myotis brandtii, Myotis lucifugus, Myotis davidii, Eptesicus fuscus, Pteropus alecto*, *Pteropus vampyrus*, and *Rousettus aegyptiacus,* and six other mammalian genomes: *Homo sapiens, Mus musculus, Bos taurus, Equus caballus, Heterocephalus glaber*, and *Monodelphis domestica*; S2 Table) using OrthoMCL software [13]. The *M. rufoniger* genes were matched and added to the orthologous clusters of *M. brandtii*’s genes, and 6,782 single-copy gene families among the 14 species were identified [7].

The divergence times of *M. rufoniger* and the 13 mammals were estimated using 1,258,141 four-fold degenerate sites from the 6,782 single-copy genes (Fig. 1). The closest clade to bats was *E. caballus* (horse) and *B. taurus* (cow) clade (diverged at 79.45 million years ago [MYA]). The *Myotis* bats diverged into its current clade approximately 11.52 MYA. *M. davidii* was estimated to be the closest to the *M. rufoniger* among the three *Myotis* bats used in this study. The divergence time between *M. rufoniger* and *M. davidii* was estimated to be 10.41 million years, indicating a fairly recent divergence which is suitable for our close species comparative genomics analyses.

**Fig 1.**
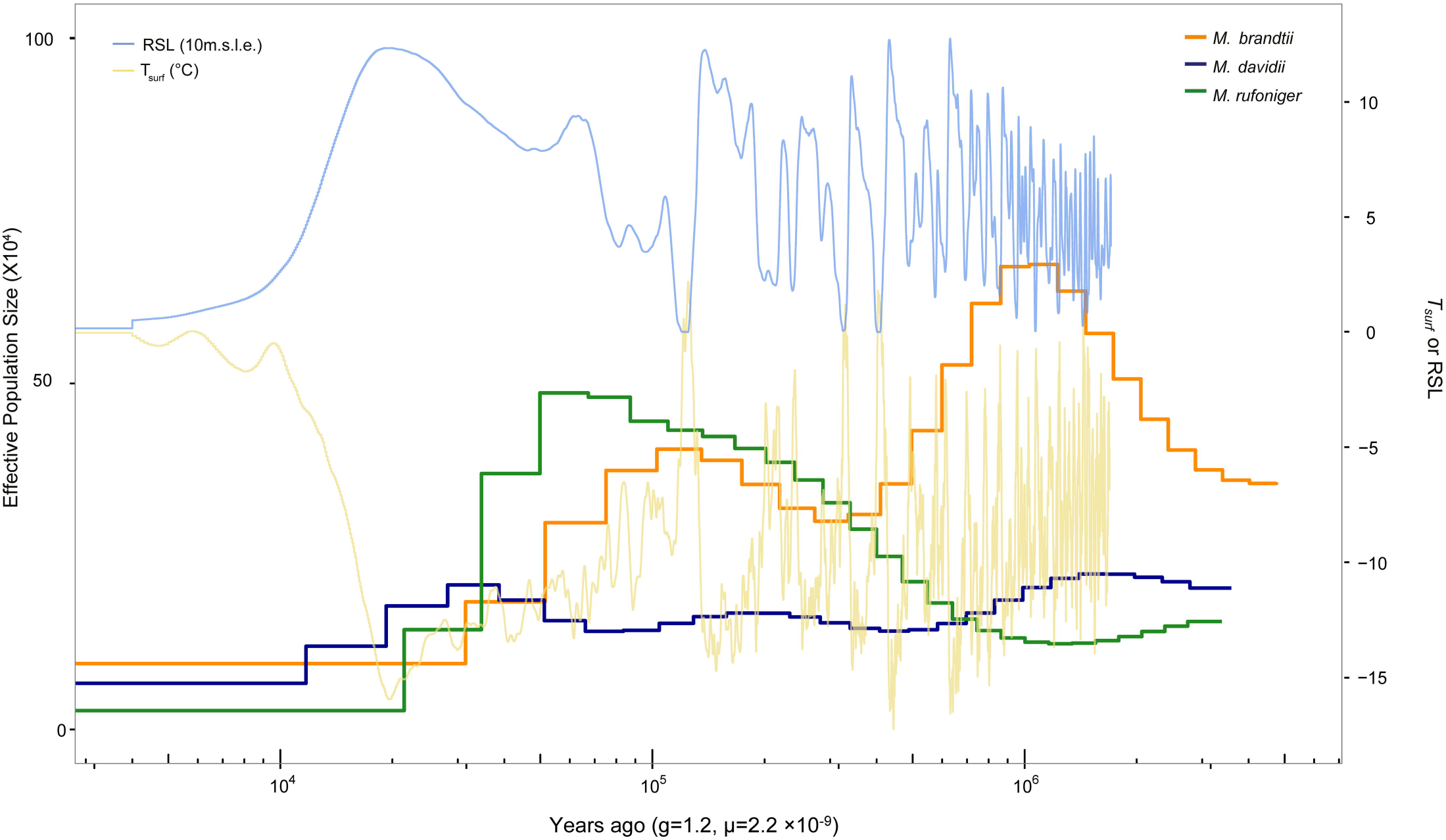
Phylogenetic relationships and divergence times in bats and mammalian species. The estimated divergence time (million years ago; MYA) is given at the nodes, with the 95% confidence intervals in parentheses. The calibration times of *M. brandtii* - *H. sapiens* (97.5 MYA), and *M. brandtii* - *P. alecto* (62.6 MYA) were derived from the TimeTree database. Colored branches and circles represent bat groups (*blue*: insect-eating microbat, *red*: fruits-eating mega bat). *M. domestica*, used as an outgroup species, was excluded in this figure.

### *M. rufoniger* specific amino acid sequences

From previous studies [3,4], bat-specific amino acid sequences within *FSHB*, *GHR*, *IGF1R*, *TP53, MDM2*, *SLC45A2*, *RGS7BP*, *RHO*, *OPN1SW*, and *CNGB3* genes were reported. They represent some general characteristics of *Myotis* bats: delayed ovulation (*FHSB*), long lifespan (*GHR* and *IGF1R*), powered flight (*TP53* and *MDM2*), echolocation (*SLC45A2* and *RGS7B*), and low vision (*RHO*, *OPN1SW*, and *CNGB3*). The same amino acid sequences within the above listed genes were also identified in the *M. rufoniger* genome (S3 Figure).

To further identify bat-specific amino acid sequences linked to environmental adaptations and unique evolutionary features, we investigated *M. rufoniger* specific amino acid changes compared to other *Myotis* bats. A total of 3,366 unique amino acid changes (uAACs) from 2,125 genes were identified in *M. rufoniger*. Among them, 1,696 uAACs from 1,228 genes were predicted to be function altering by the protein variation effect analyzer (PROVEAN) software (variant score ≤ -2.5; S8 Table) [14]. These function-altering variants and associated genes can provide insights onto *M. rufoniger* specific evolutionary adaptation. When gap-containing genes were excluded, 1,239 uAACs from 929 genes were predicted to be function altering (S9 Table). A functional enrichment analysis of the 929 genes having function-altering uAACs was performed using DAVID (Database for Annotation Visualization and Integrated Discovery) tool [15]. The probably function-altered genes were significantly enriched in reproduction related terms (Gene Ontology [GO] analysis with a *P*–value ≤ 0.05 by EASE scores [modified Fisher’s exact test] and with a 10% of false discovery rate [FDR]) including reproductive processes in a multicellular organism (*P*-value: 0.00026, 39 genes, GO: 0048609), ovulation cycle (*P*-value: 0.00052, 11 genes, GO: 0042698), and gamete generation (*P*-value: 0.0017, 31 genes, GO: 0007276; S10 Table). In *M. rufoniger*, pigment related terms were significantly enriched in the function-altered genes as in the melanin biosynthetic process (*P*-value: 0.0037, 4 genes, GO: 0042438; S10 Table). The genes are *DCT* (Dopachrome tautomerase), *SLC45A2* (Solute carrier family 45), *TYRP1* (Tyrosinase-related protein 1), and *OCA2* (Oculocutaneous albinism II). We could identify some uAACs in *DCT*, *SLC45A2*, *TYRP1*, and *OCA2* genes in the other *Myotis* bats as function-altering using PROVEAN software (Table 2; S11 Table) [14] indicating those genes are not only specific to *M. rufoniger’*s red fur color. The multiple sequence alignments and specific amino acid sequences are presented in S4 Figure.

**Table 2.** *Myotis* bats’ uAACs within melanin associated genes.

| Gene | Description | The number of uAACs<br>(The number of uAACs with PROVEAN score $\leq -2.5$) | | | |
| --- | --- | --- | --- | --- | --- |
|  |  | *M. rufoniger* | *M. brandtii* | *M. davidii* | *M. lucifugus* |
| *DCT* | Dopachrome tautomerase | 2 (2) | 0 (0) | 0 (0) | 1 (0) |
| *SLC45A2* | Solute carrier family 45 | 3 (1) | 0 (0) | 0 (0) | 0 (0) |
| *TYRP1* | Tyrosinase-related protein 1 | 2 (1) | 0 (0) | 1 (1) | 0 (0) |
| *OCA2* | Oculocutaneous albinism II | 3 (3) | 1 (0) | 0 (0) | 2 (2) |

As we were interested in the variants’ whole gene function, all the PROVEAN variant scores of each gene were summed up. We then ranked the sums of the function-altered gene candidates (S12 Table). For the top 20 genes, the same analysis was carried out in the other *Myotis* bat species for comparison (Table 3; S13, S14 Table). We ranked *M. rufoniger*’s genes according to the numbers of variants and summed variant scores, and compared them to those of other *Myotis* bat genes. From the result, *N6AMT1* (N-6 Adenine-Specific DNA Methyltransferase 1) showed the fourth-lowest sum of variants scores (sum of PROVEAN variant scores: -40.285 and number of variants: six *in M. rufoniger*; no uAAC in the other *Myotis* bats), indicating a high degree of function alteration of *N6AMT1* in *M. rufoniger*. The multiple sequence alignments and specific amino acid sequences are presented in S5 Figure.

**Table 3.**
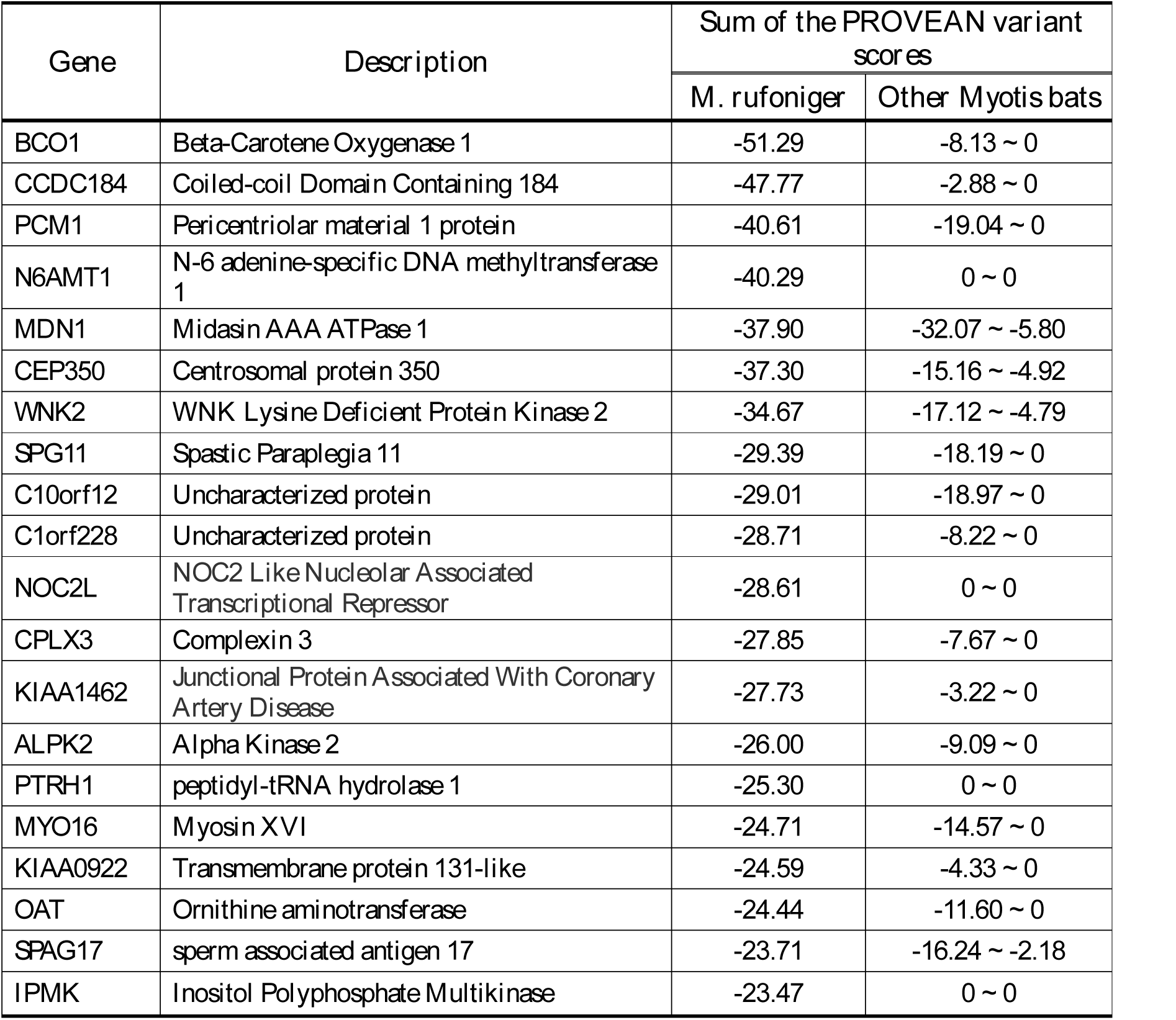
The summed PROVEAN variant score for top 20 ranked genes. The lower the more significant.

### Demographic history and the genetic diversity of *Myotis* bats

Deeply sequenced genomes allow the estimation of population structure history [10,11]. To investigate the demographic history of *Myotis* bats, a pairwise sequentially Markovian coalescent (PSMC) model inference analysis was conducted [11]. *M. rufoniger* was estimated as the most flourished species around 30 ∼ 300 k years ago compared to the other *Myotis* bats, and its demographic history showed that its population peak was 50 k years ago (Fig. 2). However, its effective population size was dramatically decreased during the last glacial period (10 ∼ 50 k years ago), and it was estimated to be the lowest in present. Also, a consistent decline in the effective population size of *Myotis* bats since ∼30 k years ago was found (Fig. 2). We further investigated the genomic diversity (which can be affected by population size) of the *M. rufoniger* and compared it to those of the other *Myotis* bats. The *M. rufoniger* genetic diversity (0.00391), based on the heterozygous SNV rate, was lower than those of the *M. davidii* (0.00471) and the *M. brandtii* (0.00614), confirming the *M. rufoniger*’s low effective population size (S13 Table).

**Fig 2.**
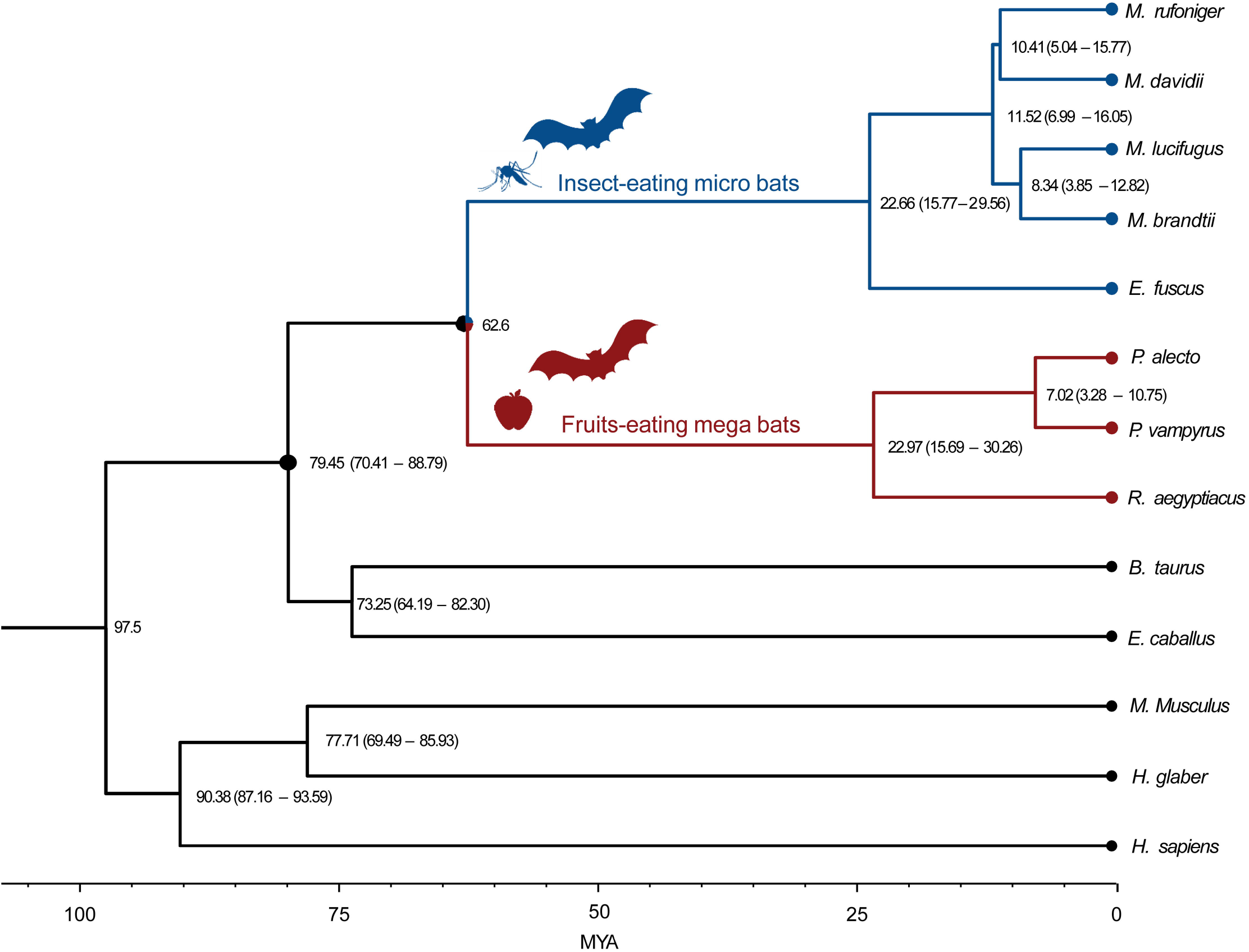
Demographic history of *Myotis* bats. *T*_surf_, atmospheric surface air temperature (T indicates temperature, surf indicates surface); RSL, relative sea level; 10 m.s.l.e., 10 m sea level equivalent; g, generation time (years); μ, mutation rate per base pair per year.

## Discussion

The function-altering variants containing genes of the *M. rufoniger* were concentrated in reproduction associated pathways. Prolonged sperm storage is a common behavior in Vespertilionidae (including genus *Myotis*, *Pipistrellus*, *Nyctalus*, *Eptesicus*, *Vesoertilio*, *Chalinolobus*, and *Plecotus*) and Rhinolophidae family species [16-18], and thus it is probably not *M. rufoniger* specific as reproduction related genes are always under strong natural selection. Therefore, such hits can be regarded as general or even a kind of artefact. Therefore, further detailed functional verification is necessary to understand the roles of each uAAC in functional categories such as reproduction.

The previous bat studies reported unique variants representing some general characteristics of *Myotis* bats: delayed ovulation (*FHSB*), long lifespan (*GHR* and *IGF1R*), powered flight (*TP53* and *MDM2*), echolocation (*SLC45A2* and *RGS7B*), and low vision (*RHO*, *OPN1SW*, and *CNGB3*) [3, 4]. We confirmed the reported amino acid sequences within the above listed genes were conserved in the *M. rufoniger* as in other bat genomes (S3 Figure), supporting that those variants are *Myotis* bat-wide not one species-specific nor individual-specific variants.

We also found that the function-altering unique variants containing the genes of the *M. rufoniger* were concentrated in a melanin associated pathway including *DCT*, *SLC45A2*, *TYRP1*, and *OCA2* genes. *TYRP1* and *DCT* genes are known to influence the quantity and the quality of melanin [19]. Mutations on the *TYRP1* gene are known to be associated with “rufous/red albinism”, causing reddish-brown skin or red hair [20-22]. The protein (melanocyte-specific transporter protein) encoded by *OCA2* gene has been reported as a transporter of tyrosine, the precursor to melanin synthesis [23]. Variants within the *SLC45A2* gene have shown association with hair color, affecting red/yellow pheomelanin pathway [24-25]. In this context, we suggest that the variants within the *DCT*, *SLC45A2*, *TYRP1*, and *OCA2* genes are likely responsible for the *M. rufoniger*’s rusty orange fur color, which distinguish it from the other bats. However, fewer but certain uAACs within the *DCT, SLC45A2*, *TYRP1*, and *OCA2* genes were also found in the other *Myotis* bats, suggesting a possible general role of those genes in bat-wide coloration. Therefore, a set of bat species with color diversity will be required for further analyses to understand the precise geno-phenotype associations in coloration.

In the *M. rufoniger* genome, *N6AMT1* gene showed a high degree of function alteration. The protein (HemK methyltransferase family member 2) encoded by *N6AMT1* gene can transform monomethylarsonous acid to dimethylarsinic acid, conferring resistance of mammalian cells to arsenic-induced toxicity [26]. An elemental analysis in the tissues from the *M. rufoniger* individual analyzed in the present study showed a very high concentration of arsenic in its intestinal tissue [27], suggesting that the *M. rufoniger* individual was exposed to food or water highly contaminated with arsenic. In this context, uAACs in *N6AMT1* can be considered as a possible chemical adaptation signature. It can be a generally stressful condition for the bats living in caves, because caves are hazardous due to various harmful elements and chemicals such as heavy metals [28]. However, until a further experimental validation is carried out, it is a very speculative hypothesis of rapid adaptation to harmful cave elements. Furthermore, this study used genomic data from only one individual for each species and the variant comparison analysis can be biased by individual-specific variants. More genomes are necessary to be sequenced to confirm that our finding’s general applicability in both *M. rufoniger* and *Myotis* bats.

The *M. rufoniger* showed the lowest effective population size compared to other *Myotis* bats from the PSMC analysis. This does not necessarily indicate that they are the most endangered as they are fairly widely spread. It is perhaps more reasonable to attribute it to very recent human encroachment. It is also shown that the population size of the *M. rufoniger* was dramatically decreased during the latter part of the last glacial period. There was a consistent decline of *Myotis* bat family’s effective population size since ∼30 k years ago. However, we cannot determine whether it is a bat-wide or *Myotis* bat specific phenomenon either, because we conducted the PSMC analysis using a small number of species. The dramatic decrease in the population size of the *M. rufoniger* could be a Korean *M. rufoniger* specific phenomenon. Therefore, a set of diverse species will be required to accurately model the bats’ effective population size history.

In conclusion, we report the first whole genome sequences of the *M. rufoniger* with bioinformatics analyses such as multiple sequence alignment, function altering uAAC prediction, functional enrichment, and population size estimation using PSMC. These provided us with some speculative insights on how bats adapted to environment sustaining the current behavioral, physiological, and demographic features such as bat’s prolonged sperm storage during reproduction, long lifespan, powered flight, low vision, echolocation, possible arsenic resistance, fur coloration, and consistently decreasing effective population sizes in their demographic history.

## Materials and Method

### Sample and genome sequencing

The partially decomposed carcass of *M. rufoniger* was found in August 2012 at Gosudonggul cave in Danyang, South Korea (coordinates: 36° 59’ 18.67“ N, 128° 22’ 53.26” E; elevation: 180.65 m) by Gosudonggul cave management office staff. The *M. rufoniger* was moved to the Natural Heritage Center of the Cultural Heritage Administration (South Korea) and kept frozen (Keeping temperature: -70□). We acquired the *M. rufoniger* sample from the Natural Heritage Center under the Cultural Heritage Administration permit in June 2013. The wing membrane tissue, which seemed not decayed, was acquired for genomic DNA sequencing (S1 Picture).

Its genomic DNA was sequenced using Illumina HiSeq2000 platform, with a read length of 100 bp and insert sizes of 566 and 574 bp from two genomic libraries. The DNA was extracted using a Tiangen Micro DNA Kit. The amount of DNA was quantified by fluorometry using a UV spectrophotometer (Tecan F200). A partial DNA degradation was identified by electrophoresis (S2 Picture), but it was judged good resulting in no significant sequencing quality problems. The genomic DNA was sheared to approximately 566 bp and 574 bp using a Covaris S2 Ultrasonicator, and then used in the preparation of whole-genome shotgun libraries according to Illumina’s library preparation protocols. The efficacy of each step of the library construction was ascertained using a 2100 Bioanalyzer (S6 Figure). The final dilutions of the two libraries were then sequenced using a HiSeq2000 sequencer with the TruSeq PE Cluster kit v3-cbot HS and TruSeq SBS kit v3-HS for 200 cycles.

### Sequencing read filtering

A DNA read was filtered out when the read’ Q20 base content was lower than 70%, using IlluQCPRLL.pl script of NGSQCToolkit (ver 2.3.3; S1 Table) [29]. We additionally downloaded genome data of the other bats (*Myotis brandtii*, *Myotis davidii*, *Myotis lucifugus*, *Pteropus Alecto*, and *Pteropus vampyrus*) from the NCBI database. Low quality DNA reads from the other bats were also filtered out using the same method (S1 Table). The possible microbial contaminated DNA reads were filtered out when the reads’ alignment scores to microbial (bacteria and fungi) genomes were higher than that to the *Myotis* bat genomes (*M. brandtii*, *M. davidii*, and *M. lucifugus*; S3 Table). We used microbial genomes that were downloaded from Ensembl database and the three *Myotis* bat genome assemblies were from NCBI. The DNA reads were mapped to the microbial and *Myotis* bat genomes using BWA-MEM algorithm of BWA (version 0.7.15) with Mark shorter split hits as a secondary option (-M) [30]. The rmdup command of SAMtools (ver 0.1.19) was used to remove PCR duplicates [31]. The possible microbial contaminated reads in the other *Myotis* bats were also filtered out by using the same method but mapping was to their own genome assemblies (S1 Table).

### Species identification

To construct the phylogenetic tree and identify the sample’s species, the mitochondrial cytochrome b consensus sequence was generated by mapping the sequencing data to the *M. rufoniger* mitochondrial cytochrome b sequence (GenBank accession number: HQ184048.1) using BWA-MEM algorithm of BWA (version 0.7.15) with Mark shorter split hits as a secondary option (–M) [30]. The reads were realigned using the GATK (version 2.5-2) RealignerTargetCreator and IndelRealigner algorithms to minimize the read alignment mismatches [32]. The rmdup command of SAMtools (ver 0.1.19) was used to remove PCR duplicates [31]. The variants from the whole genome sequencing data against the reference were called using mpileup command of SAMtools (version 0.1.19) with –E option to minimize the noise, -A option to use regardless of insert size constraint, -q 20 to consider only high quality mapping reads, and –Q 30 to consider only high quality bases [31]. The called variants were filtered using vcfutils.pl varFilter command of SAMtools (ver 0.1.19) with -d 8 option (minimal depth of 8) [31]. SNVs detected from the whole genome sequencing data against the reference were substituted to construct mitochondrial cytochrome b consensus sequence of the *M. rufoniger* using vcfutils.pl vcf2fq command of SAMtools (ver 0.1.19) with -l 5 option (no indel present within a 5 bp window) [31]. A multiple sequence alignment of the mitochondrial cytochrome b sequences was conducted using MUSCLE (version 3.8.31) program with default options [2,33]. A phylogenetic analysis was conducted using MEGA 7.0 program [34]. The phylogenetic tree was inferred by using a Maximum Likelihood method based on the Tamura-Nei model with 1,000 replicates bootstrapping [35,36]

### Genome size estimation

For a *K*-mer analysis, KmerFreq_HA command of SOAPec program in SOAPdenovo2 package was used with a *K*□= □17 option [37]. The genome size was estimated from the number of *K*-mers (depth > 3) divided by the peak depth of the *K*-mer graph.

### Mapping sequence data

The filtered sequenced reads were mapped to the three *Myotis* bat genome assemblies (*M. brandtii*, *M. davidii*, and *M. lucifugus*) using BWA-MEM algorithm of BWA (version 0.7.15) with Mark shorter split hits as a secondary option (-M) [30]. The reads were realigned using the GATK (version 2.5-2) RealignerTargetCreator and IndelRealigner algorithms to minimize the read alignment mismatches [32]. The rmdup command of SAMtools (ver 0.1.19) was used to remove PCR duplicates [31]. The variants of *M. rufoniger* were called using mpileup command of SAMtools with –E, -q 20, -A, and –Q 30 options [31]. The called variants were filtered using vcfutils.pl varFilter command of SAMtools (ver 0.1.19) with -d 8 and -D 250 options (minimal depth of 8 and maximal depth of 250) [31]. The DNA consensus sequences of *M. rufoniger* were generated by substituting the *M. rufoniger* SNVs detected from the whole genome sequencing data to each reference bat genome using the vcfutils.pl vcf2fq command of SAMtools (ver 0.1.19) with -l 5 option [31].

### Repeat annotation

RepeatMasker (version 4.0.5) was used to identify transposable elements by aligning the *M. rufoniger* consensus genome sequence against a library of *M. brandtii* with default option [38]. For the comparison of the *M. rufoniger*’s repeat annotation with the other species, the same method was used for other genome references.

### Sex determination

Male *M. davidii* reads were mapped to the *M. brandtii* (a male sample) genome assembly using BWA-MEM algorithm of BWA 0.7.15 version [30] with –M option. The reads were realigned using the GATK (version 2.5-2) RealignerTargetCreator and IndelRealigner algorithms to minimize the read mismatches [32]. The rmdup command of SAMtools (ver 0.1.19) was used to remove PCR duplicates [31]. Variants were called using mpileup command of SAMtools with –E, -q 20, -A, and –Q 30 options [31]. The called variants were filtered using vcfutils.pl varFilter command of SAMtools (ver 0.1.19) with -d 8 and -D 250 options [31]. To identify X-chromosomal scaffolds from the *M. brandtii* genome assembly, we conducted BLASTn (Blast 2.2.26) with an E-value cutoff of 1E-6 using X-chromosomal CDS sequences of human as queries [39]. Scaffolds having greater than or equal to 10 top hits were considered as X-chromosomal scaffolds.

### Comparative evolutionary analysis

Orthologous gene families between *M. rufoniger*’s sequence and 13 mammalian genomes (Seven bat genomes: *M. brandtii, M. lucifugus, M. davidii, E. fuscus, P. alecto*, *P. vampyrus*, and *R. aegyptiacus*, and six other mammals: *H. sapiens, M. musculus, B. taurus, E. caballus, H. glaber*, and *M. domestica;* S2 Table) were identified using OrthoMCL software (version 2.0.9) by matching and adding the *M. rufoniger* genes to orthologous clusters of *M. brandtii*’s genes, and 6,782 single-copy gene families among the 14 species were collected [7,13]. *M. brandtii, M. lucifugus, M. davidii, E. fuscus, P. alecto*, *P. vampyrus*, *R. aegyptiacus, H. sapiens, M. musculus, B. taurus, E. caballus, H. glaber*, and *M. domestica* genomes and gene sets were downloaded from the NCBI database (S3 Table).

To construct a phylogenetic tree and estimate the divergence time of bats and other mammals, 1,258,141 four-fold degenerate sites from the 6,782 single-copy gene families were used to conduct the multiple sequence alignment (MSA) using MUSCLE (version 3.8.31) program [33]. The phylogenetic tree was constructed among the 14 species using a Randomized Axelerated Maximum Likelihood (RAxML) program (version 8.2) [40]. In RAxML program, GTR gamma model was used as the nucleotide substitution model. In order to check the branch reliability, 1,000 rapid bootstrapping was used. The *M. domestica* was used as an outgroup species. Based on this phylogenetic tree topology, the divergence time was estimated using Reltime-ML in MEGA 7.0 program [34]. In this process, the divergence time at the node between *M. brandtii* – *H. sapiens* was constrained to be 97.5 MYA, and *M. brandtii* – *P. alecto* was constrained to be 62.6 MYA based on the TimeTree database [41].

Amino acid changes were identified by constructing MSAs among the 6,782 single copy gene families using Clustal Omega (version 1.2.4) [42]. Function-altering amino acid changes were predicted using PROVEAN (version 1.1.5) [14]. The MSAs of the *DCT*, *SLC45A2*, *TYRP1*, OCA2, and *N6AMT1* genes were manually checked, and uAACs within misaligned regions were excluded in the variant score analysis. Only homozygous amino acid variants were considered in the amino acid sequence comparison analysis to reduce bias from individual-specific variants.

### Demographic history and genetic diversity

The demographic history of the *Myotis* bats was estimated using a pairwise sequential Markovian coalescent (PSMC) program [11]. We mapped the downloaded genome sequencing data of *Myotis* bats to the *M. lucifugus* genome assembly using BWA-MEM algorithm of BWA 0.7.15 version [30] with –M option. The reads were realigned using the GATK (version 2.5-2) RealignerTargetCreator and IndelRealigner algorithms to minimize the read mismatches [32]. The rmdup command of SAMtools (ver 0.1.19) was used to remove PCR duplicates [31]. SAMtools was used to extract diploid genome information from the BAM files [31]. Options of -N25 -t15 -r -p “4+25*2+4+6” were used for the PSMC analysis. The generation time for the *Myotis* bats was estimated as sum of average maturation and gestation time (S16 Table) [43,44]. The mutation rate of *Myotis* bats was estimated by multiplying reported neutral mutation rate for mammals (2.2 × 10^−9^ per base pair per year) by the generation time (1.2 year) [45]. Atmospheric surface air temperature and global relative sea level data of the past three million years were obtained and used for this analysis [46]. The genomic diversity was calculated by dividing the number of heterozygous SNVs by its genome size (bp) [8]. For the genomic diversity calculation, variants were called using mpileup command of SAMtools with –E, -q 20, -A, and –Q 30 options [31]. The called variants were filtered using vcfutils.pl varFilter command of SAMtools (ver 0.1.19) with -d 8 and -D 250 options [31].

## Acknowledgements

Korea Institute of Science and Technology Information (KISTI) provided us with Korea Research Environment Open NETwork (KREONET) which is an academic internet connection service.

## Supporting information

**S1 Picture. *M. rufoniger* carcass picture.**

**S2 Picture. DNA sample electrophoresis.**

**S1 Figure. Phylogenetic relationship of the *M. rufoniger* sample with other bats.**

The phylogenetic relationship of *Myotis* bats was inferred from the alignment of mitochondrial cytochrome b coding sequences. The percentage of trees in which the associated taxa clustered together is shown next to the branches. Each node has its species name and GenBank accession number.

**S2 Figure. Estimation of genome size using a *K*-mer (17-mers) analysis.**

The x-axis represents depth, and the y-axis represents proportion, as calculated by the frequency at a given depth divided by the total frequency at all depths.

**S3 Figure. Previously reported bats’ unique amino acid changes.**

Previously reported bats’ unique amino acid sequence changes within *FSHB*, *GHR*, *IGF1R*, *TP53,* and *MDM2* are highlighted in yellow; (A), Alignment of *FSHB* encoded peptide sequences; (B), Alignment of *GHR*-encoded peptide sequences; (C), Alignment of *IGF1R* encoded peptide sequences; (D), Alignment of *TP53*-encoded peptide sequences; (E), Alignment of *MDM2* encoded peptide sequences; (F), Alignment of *SLC45A2*-encoded peptide sequences; (G), Alignment of *RGS7BP*-encoded peptide sequences; (H), Alignment of *RHO*-encoded peptide sequences; (I), Alignment of *OPN1SW*-encoded peptide sequences; (J), Alignment of *CNGB3*-encoded peptide sequences.

**S4 Figure. *Myotis* bats’ uAACs within *DCT*, *SLC45A2*, *TYRP1*, and *OCA2* genes.**

*Myotis* bats’ uAACs within *DCT*, *SLC45A2*, *TYRP1*, and *OCA2* genes are highlighted (yellow if PROVEAN score of variant ≤ -2.5, unless green). The domain region of human sequences are shaded in gray; (A) Alignment of *DCT* encoded peptide sequences; (B) Alignment of *SLC45A2* encoded peptide sequences; (C) Alignment of *TYRP1* encoded peptide sequences; (D) Alignment of *OCA2* encoded peptide sequences.

**S5 Figure. *M. rufoniger* specific amino acid sequence changes**

*M. rufoniger* specific amino acid sequence changes within *N6AMT1* gene are highlighted in yellow.

**S6 Figure. Sequencing library 574bp QC.**

(A) Sequencing library 574bp QC; (B) Sequencing library 566bp QC.

**S1 Table. Sequencing read filtering result.**

**S2 Table. 17-mer statistics information.**

**S3 Table. Reference genomes used in this study.**

**S4 Table. Base composition of *M. rufoniger* genome.**

The GC contents were calculated by dividing the number of [(C+G+S) – (M+K+R+Y)] ∼ [(C+G+S) + (M+K+R+Y)] by the number of (sum - gap); The proportion of A, C, G, and T, and the proportion of W, S, M, K, and Y were calculated by dividing each number by the number of (sum-gap)

**S5 Table. GC content statistics of *M. rufoniger*.**

**S6 Table. Transposon-derived repeats in the *M. rufoniger* genome.**

**S7 Table. Comparison of X-chromosomal scaffold heterozygosity.**

**S8 Table. *M. rufoniger* unique amino acid changes.**

**S9 Table. *M. rufoniger* unique amino acid changes from gap filtered sequences.**

**S10 Table. Functional enrichment of *M. rufoniger* uAACs containing genes.**

**S11 Table. PROVEAN scores of *DCT, SLC45A2*, *TYRP1*, and *OCA2* genes.**

**S12 Table. Sum of the PROVEAN scores**

**S13 Table. PROVEAN scores of top 20 ranked genes.**

**S14 Table. Sum of the PROVEAN scores of top 20 ranked genes.**

**S15 Table. Variants statistics regarding mapping of *Myotis* bat raw reads to the *M. lucifugus* reference**

**S16 Table. Generation time calculation.**

